# Anhedonia severity mediates the relationship between attentional networks recruitment and emotional blunting during music listening

**DOI:** 10.1101/2023.12.21.572579

**Authors:** Marie-Stephanie Cahart, Vincent Giampietro, Laura Naysmith, Fernando Zelaya, Steven Williams, Owen O’Daly

## Abstract

Recent studies have reported atypical emotional processing in individuals with greater levels of anhedonic depressive symptoms. However, the relationship between brain networks dynamics and moment-to-moment affective responses to naturalistic paradigms, as emotions are unfolding, remains unclear. In this study, we used the unique temporal characteristics of music to investigate behavioural and brain network dynamics as a function of anhedonic depressive symptoms severity in healthy adults during an emotionally provocative music listening task. Thirty-one neurotypical participants aged 18-30 years were required to continuously rate happy, neutral and sad pieces of music whilst undergoing MRI scanning. They were also asked to fill in questionnaires assessing their levels of anhedonic depressive symptoms. Using a novel fMRI analysis method called Leading Eigenvector Dynamics Analysis (LEiDA), we found an increased probability of occurrence of attentional networks and a blunted emotional response to both happy and sad pieces of music in participants with greater levels of anhedonic depressive symptoms. More specifically, anhedonic depressive symptoms mediated the relationship between attentional networks recruitment and emotional blunting. Furthermore, the elevated recruitment of attentional networks during emotional pieces of music carried over into subsequent neutral music. Future studies are needed to investigate whether these findings could be generalised to a clinical population (i.e., Major Depressive Disorder).

## Introduction

Characterized by diminished pleasure or interest and significant impairments in daily functioning and quality of life, depressive disorders are among the most common affective disorders and are considered the largest contributors to global disability (World Health Organization, 2017). In particular, major depressive disorder (MDD) is regarded as the most severe form of depressive disorders, with atypical emotional reactivity to positive and negative stimuli shown to represent one of the core impairments of the illness (see Bylsma et al., 2008 for a meta-analysis). It is worth noting, however, that depressive symptoms are common and vary significantly in severity both in clinical (Thapar et al., 2022) and non-clinical populations (Sütterlin et al., 2012). In fact, in the general population, individual variation in the way and degree to which individuals react to emotional stimuli around them has been strongly associated with differences in mental illness and suicidal ideation later in life (Polanco-Roman et al., 2018). However, to date, the nature of the observed effects (i.e., increased or reduced reactivity to positive and negative cues) remains unclear, especially regarding negative emotional reactivity (Saxena et al., 2017).

At present, there are three main perspectives on emotional reactivity in depression, characterised by (i) an attenuated response to positively valenced stimuli (i.e., Positive Attenuation theory), (ii) a potentiated reactivity to negatively valenced cues (i.e., Negative Potentiation theory), or (iii) a reduced reactivity to both positive and negative information (i.e., Emotion Context Insensitivity; ECI) (Rottenberg et al., 2005). In a large meta-analysis covering studies which used a range of emotional stimuli such as images, reward tasks and stress tasks, Bylsma et al. (2008) found evidence for a blunted emotional response to both positive and negative stimuli in MDD, providing clear support for the ECI theory (Rottenberg et al., 2005). The ECI phenomenon has also been endorsed by more recent studies using valence-loaded images in MDD (Vanderhasselt et al., 2016), as well as in response to sad music in children with greater levels of depressive symptoms, drawn from the general population (Somers et al., 2020). In contrast, Vuoskoski and Eerola (2011) did find a negative correlation between happiness ratings and anger, fatigue and depression scores in the context of music listening. However, they also reported that neuroticism, anger, depression and tension scores were positively correlated with perceived sadness, a finding conforming with the Negative Potentiation theory, rather than with the ECI phenomenon. Liljeström, Juslin and Västfjäll (2013) also found that neuroticism scores were negatively associated with enjoyment-pleasure ratings, but positively correlated with sadness, anger and anxiety ratings, which also aligns with the findings of Ladinig and Schellenberg (2012) who also observed a positive relationship between neuroticism and felt sadness.

It has been argued that these discrepancies across studies could be explained by the fact that most emotion studies, until recently, have used methods where participants are required to provide a single subjective rating for a given emotional cue (Sonkusare et al., 2019). This method has been questioned, as it does not allow for the study of emotional states as they unfold over time (Sonkusare et al., 2019). We argue that gaining a better understanding of the interindividual variation in the time-varying behavioural and neural correlates of emotional reactivity in the broader population may, ultimately, lead to identifying markers of vulnerability to affective disorders that could inform early prevention and intervention strategies (Noyes et al., 2022).

In fact, a growing number of studies have shown that psychological well-being is as reflective of the variability of how one feels over time, as it is of how they feel on average (see Houben et al., 2015 for a meta-analysis). This insight has led to the development of two dynamic summary measures of subjective affective variability over time (Houben et al., 2015; Trull et al., 2015; Wichers et al., 2015). First, the within-subject standard deviation (STD) provides information about the variance of the participant’s emotional states over a given time period (Eid & Diener, 1999). Here, high and low values reflect heightened emotional reactivity and blunted responses, respectively. Second, the within-subject root mean square of the successive differences (RMSSD; Ebner-Priemer & Trull, 2009) reflects moment-to-moment fluctuations in emotional reactivity. Here, higher scores reflect greater emotional lability, characterised by more significant emotional swings from one moment to the next, while lower scores suggest a less flexible emotional response. Overall, the consensus deriving from Houben et al. (2015)’s meta-analysis was that increased STD and RMSSD were associated with lower psychological well-being. However, it is worth noting that these studies employed experience sampling or diary methods (ESM), whereby participants were invited to rate how they felt every few hours throughout the day in the absence of any standardized stimulus, which makes it difficult to make formal comparisons across individuals in terms of their emotional reactivity to specific valence-loaded cues.

In the last decade, neuroimaging studies have provided considerable insight into the neural basis of emotional reactivity as a function of emotional wellbeing in the context of music listening. Using a region of interest analysis, Park et al. (2013) found positive correlations between neuroticism severity and activation in the orbitofrontal cortex, the basal ganglia and the insula in response to happy music. However, their sample size was very small (i.e., 12 participants), and the pieces of music used as a reference had previously been validated to convey a pleasant rather than neutral state, potentially triggering some affective responses. Additionally, the participants were not required to rate how each piece of music made them feel. Instead, they were simply asked to passively listen to the music, making it difficult to establish whether the emotional induction worked. Trait anhedonia, the inability to experience pleasure from typically pleasant stimuli and life events, has also been shown to negatively correlate with pleasantness ratings and with activity in the nucleus accumbens, the basal forebrain, the hypothalamus, the anterior insula and the orbitofrontal cortex in a music listening task in healthy adults (Keller et al., 2013). However, as in most music studies, the subjective ratings here were collected following the scan, using a single rating for each piece of music, which again precludes the examination of the dynamics of emotional reactivity.

In this study of healthy adults, we used the unique temporal characteristics of music to explore the dynamic features of continuous subjective emotional experience in response to happy, neutral and sad emotional pieces of music as a function of anhedonic depressive symptoms severity. We combined continuous reporting measures with a novel fMRI analysis method called Leading Eigenvector Dynamics Analysis (LEiDA; Cabral et al., 2017). LEiDA affords the exploration of dynamic neural processes at the group level while preserving idiosyncratic information about moment-to-moment BOLD fluctuations. Using this approach, greater subclinical depression scores have previously been associated with increased activity of the Default Mode Network (DMN) and fronto-parietal networks, whereas visual and Dorsal Attention networks were diminished in their prominence (Alonso-Martínez et al., 2020). However, those findings were observed using resting-state fMRI. In fact, even though resting-state fMRI is easy to collect and has helped shed light on the functional architecture of the brain, recent studies have shown that it is outperformed by naturalistic paradigms when it comes to predicting behaviour (Finn & Bandettini, 2021; Greene, Gao, Scheinost & Constable, 2018). Resting-state has also been linked to lower test-retest reliability of functional connectivity metrics (Wang et al., 2017; Zhang et al., 2022), as well as a difficulty inferring specific mental processes from unconstrained brain activity (Finn, 2021; Poldrack, 2006), hence why we decided to focus on rich naturalistic paradigms such as music in this study.

Based on the findings observed in previous emotion and LEiDA studies described above, we hypothesized that we would observe:

- An atypical pattern of emotion dynamics in healthy participants with higher levels of anhedonic depressive symptoms, as evidenced by a significant correlation between anhedonic depressive scores and the emotional STD and RMSSD metrics.
- An atypical behaviour of the DMN, fronto-parietal, visual and Dorsal Attention networks in healthy participants with higher levels of anhedonic depressive symptoms, as evidenced by a significant correlation between anhedonic depressive scores and LEiDA metrics (i.e., probability of occurrence and mean lifetime) for each network.
- An atypical behaviour of the DMN, fronto-parietal, visual and Dorsal Attention networks in participants with an atypical pattern of emotion dynamics, as evidenced by a significant correlation between emotion dynamics metrics (STD and RMSSD) and LEiDA metrics (i.e., probability of occurrence and mean lifetime) for each network.

## Materials and methods

### Participants

Thirty-nine neurotypical right-handed adults initially participated in the study after providing written informed consent. All participants were recruited based on the following inclusion criteria: between 18 and 30 years of age, good physical health, absence of any psychiatric or neurological disorder, and absence of any MRI counter-indications (i.e., pacemaker, metal in the body, claustrophobia). Of the 39 participants who took part, one had incomplete fMRI data and two reported feeling excessively distracted by the background noise generated by the scanner, resulting in an inability to feel emotions that matched those evoked by the songs; these participants were therefore discarded from the analyses. A further three participants had excessive head motion (> 3.3mm), and another two had more than 15% invalid scans as determined by the CONN toolbox Version 20b (Whitfield-Gabrieli & Nieto-Castanon, 2012). These participants were also excluded. Consequently, the final number of participants included in the analyses was 31 (17 males and 14 females) (M=22.6±4.0 years). The study was approved by the King’s College London Human Research Ethics Committee (number HR-19/20-18771). After data acquisition was completed, the researchers visually inspected the scans to check for artifacts, and a qualified neuroradiologist also reviewed the images to rule out any major neural anomaly, in line with the policies of King’s College London’s Department of Neuroimaging.

### MRI data acquisition

Participants were all scanned in the same 3-Tesla MR scanner (Discovery MR750, General Electric, Milwaukee, WI, USA) at the Centre for Neuroimaging Sciences (Institute of Psychiatry, Psychology and Neuroscience; King’s College London). The MRI data was collected by experienced radiographers using a 12-channel head coil. Anatomical T1-weighted images (MPRAGE) had the following parameters: TR=7.35ms, TE=3.04ms, flip angle=11°, slice thickness=1.2mm, in-plane resolution 1.05mm. The functional images were collected using a 2D multi-slice, gradient-recalled Echo-Planar Imaging (EPI) sequence with the following parameters: TR=2s, TE=33ms, flip angle=75°, slice thickness=3mm, field of view=240mm, 64x64 matrix. The total duration of the fMRI run was 8 minutes and 27 seconds. Four dummy scans were acquired at the beginning of the time series, to allow the signal to achieve steady state. Those four volumes were not part of the analysed time series. All participants were provided with earplugs and padded headphones to limit any discomfort deriving from the background noise generated by the scanner.

### MRI data pre-processing

The data were pre-processed using the CONN toolbox Version 20b (Whitfield-Gabrieli & Nieto-Castanon, 2012), MATLAB R2020a (MathWorks, Natick, MA, USA). The pre-processing consisted of realignment to correct for volume-to-volume head motion, co-registration of the functional data to the anatomical image, and spatial normalization into the Montreal Neurological Institute (MNI) standardized space by means of unified segmentation of the T1 weighted structural image. Finally, the normalized functional MRI data were smoothed with full width at half-maximum isotropic Gaussian kernel of 8 mm. The artifact rejection tool (ART), implemented in CONN (http://www.nitrc.org/projects/artifact_detect), was used to detect outlier volumes in the timeseries with respect to head motion and global signal changes. One covariate for each outlier volume was then entered in the denoising step to lessen the contribution of those scans on the results of the fMRI analyses. The anatomical component-based noise correction approach (aCompCor; Behzadi et al., 2007) was also used in order to ensure that physiological and other sources of noise which are unlikely to be neuronal in origin were regressed out.

### Stimuli and procedures

During the fMRI acquisition, all participants were required to listen to 13 pieces of classical music and asked to continuously rate how each piece of music made them feel on a scale of -6 (very sad) to +6 (very happy). These pieces of music were chosen as they have previously been identified as eliciting happy, neutral and sad emotional states in healthy adults (Mitterschiffthaler et al., 2007). The participants listened to the auditory stimuli through headphones, and they responded by using a two-button box. Pressing the left button made the slider move left towards -6, while pressing the right button made it move right towards +6. The equipment was tested before the scan started to ensure auditory clarity. All songs were presented in the exact same order for all participants to facilitate analysis methods, with 2 seconds of silence separating each pair of pieces of music. Overall, the run consisted of 3 happy songs, 3 sad songs and 7 neutral songs. It started with a neutral song, and then each sad song and each happy song was followed by a neutral song. The name of each musical piece, the name of the composer, the order of presentation, the song type and the start and end times are presented in Table 1. The pieces of music slightly varied in length to minimise abrupt and harsh stops.

**Table 1.** Names of each music piece, order of presentation, start and end times, composer and song types.

| Order of presentation | Start time (in secs) | End time (in secs) | Musical pieces | Composer | Song type |
| --- | --- | --- | --- | --- | --- |
| 1 | 0 | 29 | Water music – passepied | Handel | Neutral |
| 2 | 31 | 65 | Kol Nidrei | Bruch | Sad |
| 3 | 67 | 109 | Violin romance no.2 in F major | Beethoven | Neutral |
| 4 | 111 | 147 | Radetzky march | Strauss | Happy |
| 5 | 149 | 183 | L’oiseau prophete | Schumann | Neutral |
| 6 | 185 | 225 | Suite for violin & orchestra in A minor | Sinding | Sad |
| 7 | 227 | 259 | Clair de lune | Debussy | Neutral |
| 8 | 261 | 301 | A little night music – allegro | Mozart | Happy |
| 9 | 303 | 337 | Water music menuet | Handel | Neutral |
| 10 | 339 | 381 | Concerto de Aranjuez – Adagio | Rodrigo | Sad |
| 11 | 383 | 425 | The Planets – Venus | Holst | Neutral |
| 12 | 427 | 467 | A little night music – Rondo Allegro | Mozart | Happy |
| 13 | 469 | 507 | La Traviata – Prelude to the 1<sup>st</sup> scene | Verdi | Neutral |

#### Self-report measures

Prior to the scan, participants were required to fill in questionnaires aimed at assessing their familiarity with each piece of music, their music background, anhedonic depression severity, neuroticism, rumination levels and trait emotional reactivity (see details below).

##### Personality and mental health questionnaires

The participants were then required to fill in the following questionnaires: the anhedonic depression subscale of the Mood and Anxiety Symptoms Questionnaire (MASQ-AD; Clark & Watson, 1991), the neuroticism subscale of the Big Five Inventory (BFI; John & Srivastava, 1999),the rumination subscale of the Rumination-Reflection Questionnaire (RRQ; Trapnell & Campbell, 1999) and the short version of the Perth Emotional Reactivity Scale (PERS-short; (Preece, Becerra & Campitello, 2019).

#### Mood and Anxiety Symptoms Questionnaire – Anhedonic Depression subscale (MASQ-AD)

The 14-item MASQ-AD subscale of the MASQ was chosen to evaluate participants’ levels of anhedonic depression (Clark & Watson, 1991). This subscale has previously been used to examine the neural basis of anhedonic depression in the context of music listening, and it was specifically chosen for this study because it has demonstrated satisfactory variance and high sensitivity to individual differences in a healthy adult population (Keller et al., 2013). It comprises items that assess loss of interest and enjoyment concerning a wide-range of everyday situations. For each item, participants were required to indicate on a five-point scale (1=not at all, 5=extremely) to what extent each statement applied to them. Scores were then summed up across all 14 items, and the overall scores ranged between 14 to 70. A score of 14 indicates an absence of anhedonic symptoms, while a score of 70 reflects high levels of anhedonic depression. The MASQ-AD has shown good internal consistency alpha in university samples (α=.80; Harra & Vargas, 2023).

For this paper, we focussed on anhedonic depressive symptoms and therefore only the MASQ-AD and key associated findings will be further described in this paper. More information about the other scales can be found in Supplementary Material 1a.

##### Familiarity ratings

The participants were first required to listen to each piece of music through the Lime Survey online tool (limesurvey.org) and rate them in terms of how familiar they were with each of them on a five-point scale (1=I have never heard this piece of music before, 5=I know this piece of music very well). The mean and standard deviation of the familiarity ratings were then calculated for each song across all participants.

##### Music background

The music background questionnaire consisted in four statements which the participants were asked to rate in terms of how much they each applied to them (1=not at all, 5=extremely): “I enjoy listening to music” (question 1), “I enjoy listening to classical music” (question 2), “I have a broad knowledge of classical music (e.g., I can recognise and name different classical music pieces, composers, instruments etc)” (question 3), and “I can play a musical instrument” (question 4).

### Behavioural analyses

For all behavioural analyses, normality of each variable was tested using the Shapiro-Wilk test (significance was set at 0.05).

#### Familiarity ratings and music background

Pearson’s correlations were carried out between music background questions and summary measures of the familiarity ratings for each song type (i.e., Familiarity_H for the familiarity with the happy songs, Familiarity_S for the familiarity with the sad songs, Familiarity_N for the familiarity with the neutral songs). Question 3 (i.e., I have a broad knowledge of classical music) consistently exhibited strongest associations with all four familiarity summary measures, therefore the question 3 scores were added to gender and age as a nuisance covariate for the subsequent analyses. Full details are provided in Supplementary Material 2a.

From this point, reference to music background/familiarity will refer to the responses to question 3.

#### MASQ-AD and mean subjective ratings

We carried out a repeated-measures ANOVA, followed by two post-hoc paired t-tests to explore whether there were significant differences between mean subjective ratings for ‘happy’, ‘neutral’ and ‘sad’ pieces of music. We then correlated mean ratings with MASQ-AD scores to explore associations between the anhedonic depression symptoms and mean subjective ratings for each song type. Full details are provided in Supplementary Material 3a.

In all cases, the threshold for significance was set at p<0.05.

#### MASQ-AD and emotional blunting/variability

For each participant, the within-subject standard deviation (STD) and the within-subject root mean square of successive differences (RMSSD) were calculated as follows, in line with Jahng, Trull and Wood (2008), where n is the number of timepoints, μis the average of the subjective ratings over the entire task, x_i_ is the subjective rating at time t, and x_i+1_ is the subjective rating at time t+1:

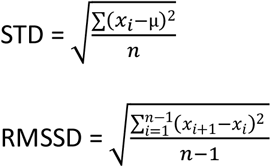

To identify whether anhedonic depression was related to either of the participants’ emotional blunting/variability metrics, we then calculated Pearson’s correlation coefficients between the MASQ-AD and the within-subject STD and RMSSD metrics, controlling for age, gender and music background/familiarity.

### LEiDA analyses

The LEiDA analyses were carried out in MATLAB R2020a (MathWorks, Natick, MA, USA) using scripts adapted from Cabral et al., 2017 (https://github.com/juanitacabral/LEiDA). N=105 regions of interest (ROI) defined anatomically based on the Harvard-Oxford cortical atlas were extracted from the CONN toolbox (Whitfield-Gabrieli et al., 2012). In line with previous work by Cabral et al. (2017), the cerebellar ROIs were excluded from the analyses because of the absence of cerebellar networks within the Yeo parcellation used for this study (Yeo et al., 2011).

Full details of the implementation of the LEiDA method can be found in Cabral et al. (2017), but in brief: for each ROI, the timeseries of the blood oxygen level dependent (BOLD) signal was first Hilbert-transformed to create an analytic signal which captures the time-varying phase of the BOLD oscillations. We then calculated the degree to which BOLD phases were synchronised between pairs of ROIs at each timepoint t, as reflected in the dynamic phase-locking matrix dPL(t) (Figure 1).

**Figure 1.**
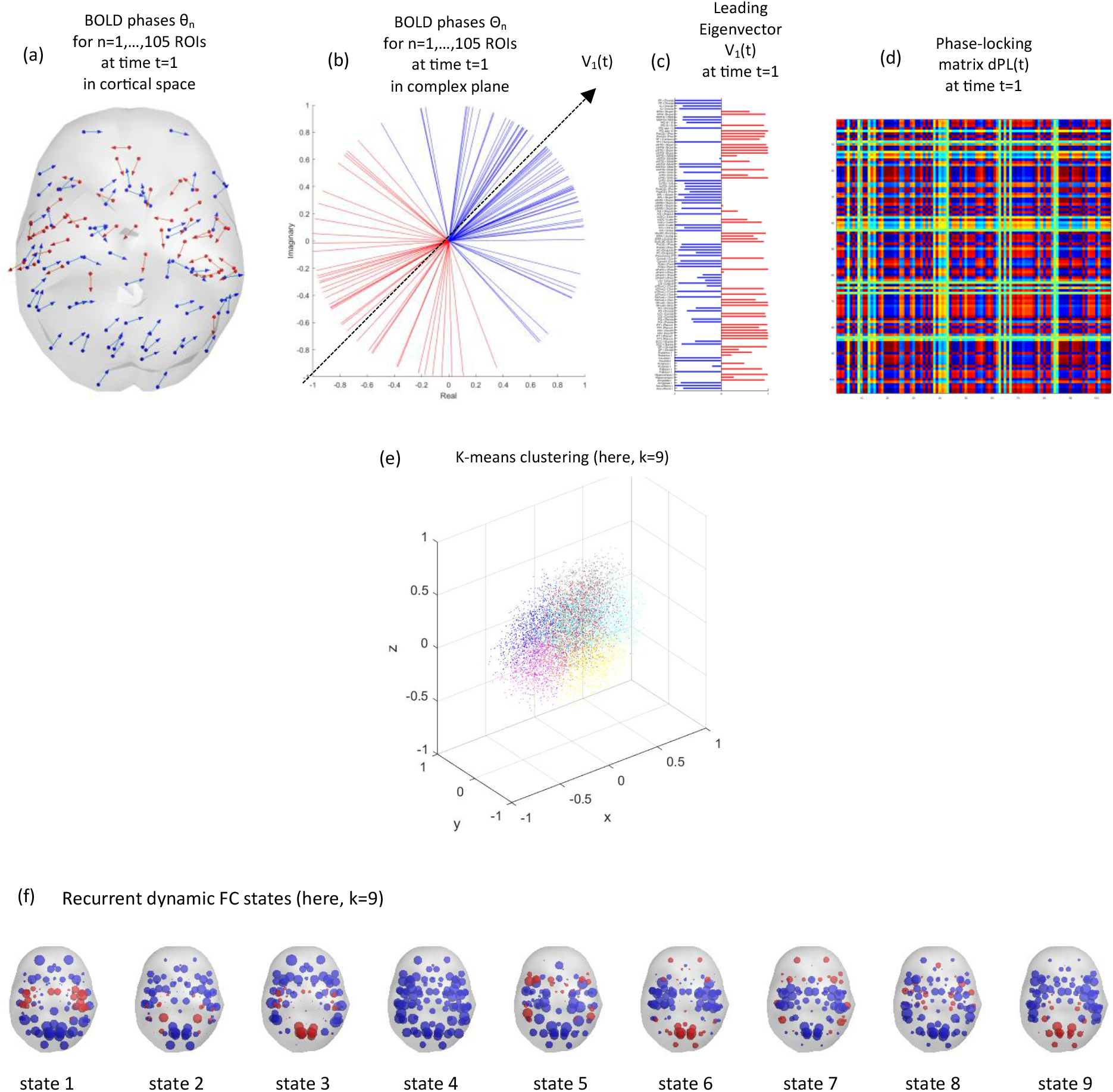
Identification of recurrent phase-locking patterns (or states) in fMRI signals. At each timepoint (here, first volume t=1), BOLD phases of each ROI are represented in (a) cortical space, where each arrow reflects the phase orientation of a given ROI and is originated from the centre of gravity of that ROI, and (b) in complex plane, where each arrow is centred at the same origin and the leading eigenvector V_1_ is represented as a dashed arrow. Phases are divided into two communities (i.e., blue or red) depending on the direction they project onto V_1_. (c) Each element in the horizontal bar plot captures the relative contribution of a given ROI to V_1_ at a given timepoint (here, first volume t=1). (d) The 105x105 dynamic phase-locking matrix dPL(t) reflects the degree of alignment, or synchrony, between pairs of ROI phases at time t (here, first volume t=1). The warmer the colour, the more synchronised the ROIs. For each participant, we obtain a leading eigenvector V_1_ at each timepoint t. (e) All leading eigenvectors V_1_ across all timepoints and all participants are then divided into k clusters using k-means clustering (here, k=9). (f) Each cluster, or state, is illustrated in cortical space. Each sphere reflects the centre of gravity of a given ROI and is coloured depending on the direction it projects onto the leading eigenvector of that state.

The next step consisted in calculating a leading eigenvector V_1_(t) for each dynamic phase-locking matrix dPL(t) to identify recurrent patterns in the dPL with reduced dimensionality. In particular, V_1_(t) contains N elements (i.e., ROIs) which each have either a positive or a negative sign depending on the direction their phase projects onto V_1_(t). When all elements of V_1_(t) have the same sign (i.e., negative), then all BOLD phases are pointing in the same direction (i.e., globally synchronised), which reflects global coherence mode. In contrast, a positive sign reflects meaningful functional networks that detach from the global coherence mode (i.e., desynchronised from the rest of the brain) and that dominate at a given timepoint (Lord et al., 2019; Vohryzek et al., 2020).

We then used k-means clustering to iteratively cluster all leading eigenvectors V_1_(t) into k=5 to k=10 network states and the Dunn score was calculated in order to detect the number of states k that best explained the data.

Once the optimal number of states k was identified, three LEiDA metrics were then calculated for each cluster or state. The probability of occurrence of a given state reflects the percentage of timepoints during which the state dominates during the scan; the lifetime represents the mean number of consecutive timepoints during which the cluster dominates; and the switching probability refers to the probability of switching from one state to another, normalised by the probability of the occurrence of the state the transition is starting from.

To assign to each state a meaningful reference label based on known functional networks, we then calculated the spatial similarities between each state and seven resting-state networks previously identified by Yeo et al. (2011). This consisted in computing the Pearson’s correlation coefficients between each of the seven networks and the centroids V_k_ previously obtained from the k-means clustering analysis, following the methodology described in Vohryzek et al. (2020). Significance was set at p<0.01/k.

For all LEiDA analyses, correction for multiple comparisons was implemented for each metric using False Discovery Rate (FDR) correction (Benjamini & Hochberg, 1995), and age, gender and music background/familiarity were controlled for.

#### Across the entire task

##### MASQ-AD and LEiDA metrics

We first investigated whether the severity of anhedonic depression symptoms was associated with each of the LEiDA measures across the entire task by calculating the Pearson’s correlation coefficients between the MASQ-AD and each LEiDA metric for each state.

##### Emotional blunting/variability and LEiDA metrics

We then explored whether each of the LEiDA metrics was associated with emotional blunting/variability by calculating the Pearson’s correlation coefficients between each LEiDA metric and the within-subject STD, and between each LEiDA metric and the within-subject RMSSD for each state.

##### Mediation analysis

To better understand the relationship between the LEiDA metrics, MASQ-AD scores and emotional blunting, we carried out a mediation analysis using IBM SPSS Statistics version 29 and macro-programme PROCESS version 4.2 (Hayes, 2013). We used the LEiDA metrics as an independent variable (IV), emotional blunting as a dependent variable (DV) and MASQ-AD as the mediator (M). A series of regression models were then fitted. The first step consisted in predicting M using IV (i.e., a-path) and then predicting DV using M (i.e., b-path). The second step consisted in measuring the indirect effect of IV on DV through M, by multiplying a and b. This allowed us to explore how M mediates the relationship between IV and DV. In order to investigate whether the mediation model was partial or full, we also estimated the direct effect of IV and DV, as well as the total effect.

#### Focussing on each song type

Previous studies have found a stronger brain/behaviour relationship during less emotional moments of a naturalistic task in depressed adolescents (Gruskin et al., 2020), and carry-over effects into subsequent conditions have also been previously observed in the context of sad emotional states (Minaeva et al., 2021) and lethargy (Ebrahimi et al., 2021) in depression. Therefore, we decided to explore further whether the relationship between the anhedonic depressive symptoms and the probability of occurrence of a given state was related to the valence of the song being played. We first extracted LEiDA metrics for each song type, only for the states where a significant correlation was observed between the MASQ-AD and their LEiDA metrics across the entire song. For each relevant state, each LEiDA metric was averaged across all songs within the same song type. We then calculated the Pearson’s correlation coefficients between the MASQ-AD and each of the LEiDA measures of a given state for each song type.

### Behavioural results

All of the behavioural variables were normally distributed, as reflected by a non-significant Shapiro-Wilk normality test (p>0.05).

### Descriptive statistics

#### Personality and mental health questionnaires

The mean and between-subject standard deviation for the MASQ-AD were 34.26 and 8.843, respectively. Descriptive statistics representing the mean and between-subject standard deviation for the other personality and mental health questionnaires can be found in Supplementary Material 1b, alongside correlations between each pair of questionnaires, and associated p-values.

We chose to focus on the MASQ-AD subscale for the rest of the analyses because the focus of this study was on anhedonic depressive (AD) symptoms, and because of the highly correlated scores between MASQ-AD and the other four questionnaires.

#### Subjective ratings and familiarity ratings

Descriptive statistics representing the mean and standard deviation of the subjective ratings for each song can be found in Table 2.

**Table 2.** Descriptive statistics providing the mean and between-subject standard deviation (SD) of the subjective ratings and familiarity ratings for each song. Participants (n=31) were required to continuously rate each song in terms of how they made them feel, from -6 (very sad) to +6 (very happy) and were also asked to indicate how familiar they were with each song (1=I have never heard this piece of music before, 5=I know this piece of music very well).

| **Musical pieces** | **Mean (SD)**<br>**subjective**<br>**ratings** | **Mean (SD)**<br>**familiarity**<br>**ratings** |
| --- | --- | --- |
| ***Happy pieces*** |  |  |
| Radetzky march | 3.92 (1.55) | 2.65 (1.43) |
| A little night music – allegro | 3.75 (1.67) | 3.10 (1.47) |
| A little night music – Rondo Allegro | 3.24 (1.60) | 2.13 (1.12) |
| ***Sad pieces*** |  |  |
| Kol Nidrei | -2.06 (2.06) | 1.48 (0.89) |
| Suite for violin & orchestra in A minor | -2.98 (1.59) | 1.39 (0.72) |
| Concerto de Aranjuez – Adagio | -2.48 (1.88) | 1.77 (1.20) |
| ***Neutral pieces*** |  |  |
| Water music – passepied | 1.28 (2.03) | 2.26 (0.96) |
| Violin romance no.2 in F major | 1.15 (2.24) | 2.29 (1.16) |
| L’oiseau prophete | -0.05 (1.50) | 1.29 (0.64) |
| Clair de lune | -0.61 (2.83) | 3.45 (1.41) |
| Water music menuet | 1.15 (1.38) | 1.77 (1.06) |
| The Planets – Venus | -1.62 (1.91) | 1.74 (1.21) |
| La Traviata – Prelude to the 1<sup>st</sup> scene | 1.49 (1.72) | 2.06 (1.09) |

### Familiarity ratings and music background

Full details about the mean and standard deviation of the music background questions and familiarity ratings for each song type can be found in Supplementary Material 2b.

Full details about the coefficients of determination and correlation coefficients between the music background questions and the familiarity scores are also provided in Supplementary Material 2b.

As question 3 consistently exhibited the highest coefficients of determination and correlation coefficients for the four familiarity summary measures, the question 3 scores were used as a nuisance covariate for the subsequent analyses.

### MASQ-AD and mean subjective ratings

Mean ratings were significantly higher happy compared to neutral songs and significantly lower for sad compared to neutral songs. Additionally, MASQ-AD significantly negatively correlated with happy ratings, and significantly positively correlated with sad ratings. Significance was set a 0.05. Full details are provided in Supplementary Material 3b.

### MASQ-AD and emotional blunting/variability

There was a large, negative, correlation between the MASQ-AD and the within-subject STD of the subjective ratings (r=-0.630, pFDR<0.001), and between the MASQ-AD and the within-subject RMSSD of the subjective ratings (r=-0.656, pFDR<0.001). An illustration is provided in figure 2a and b.

**Figure 2.**
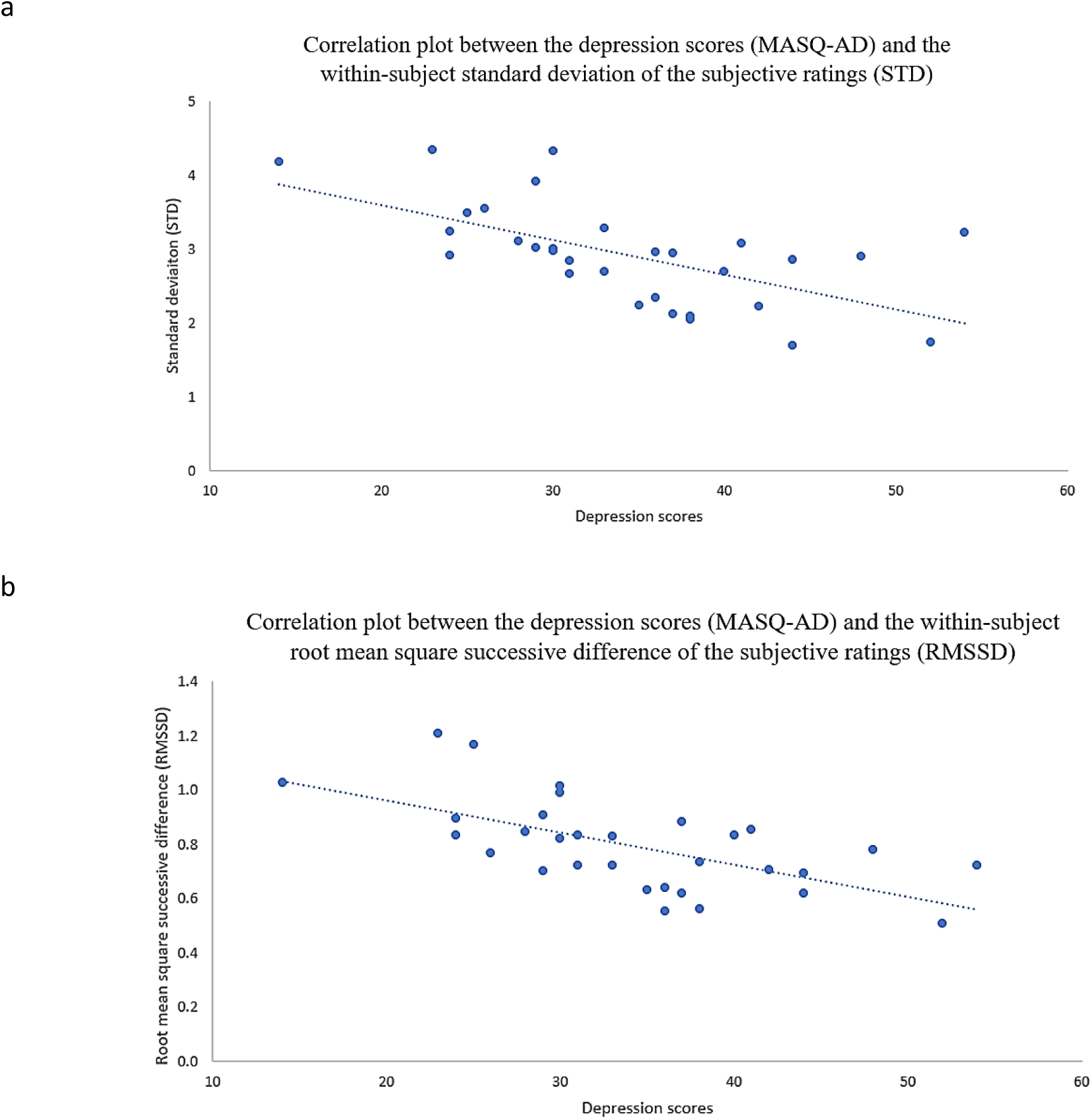
Scatter plots representing the depression scores (MASQ-AD; x axis) and (a) the within-subject standard deviation of the subjective ratings (STD; y axis) and (b) the within-subject root mean square successive difference of the subjective ratings (RMSSD; y axis), with trendlines.

Figure 3 illustrates the lower amplitude (i.e., lower STD) in subjective ratings displayed by participants whose MASQ-AD scores fell into the top 33% of the range of MASQ-AD values (i.e., (a) greater levels of anhedonic depressive symptoms) compared to those in the middle 33% (i.e., (b)), and those in the bottom 33% (i.e., (c) lower levels of anhedonic depressive symptoms).

**Figure 3.**
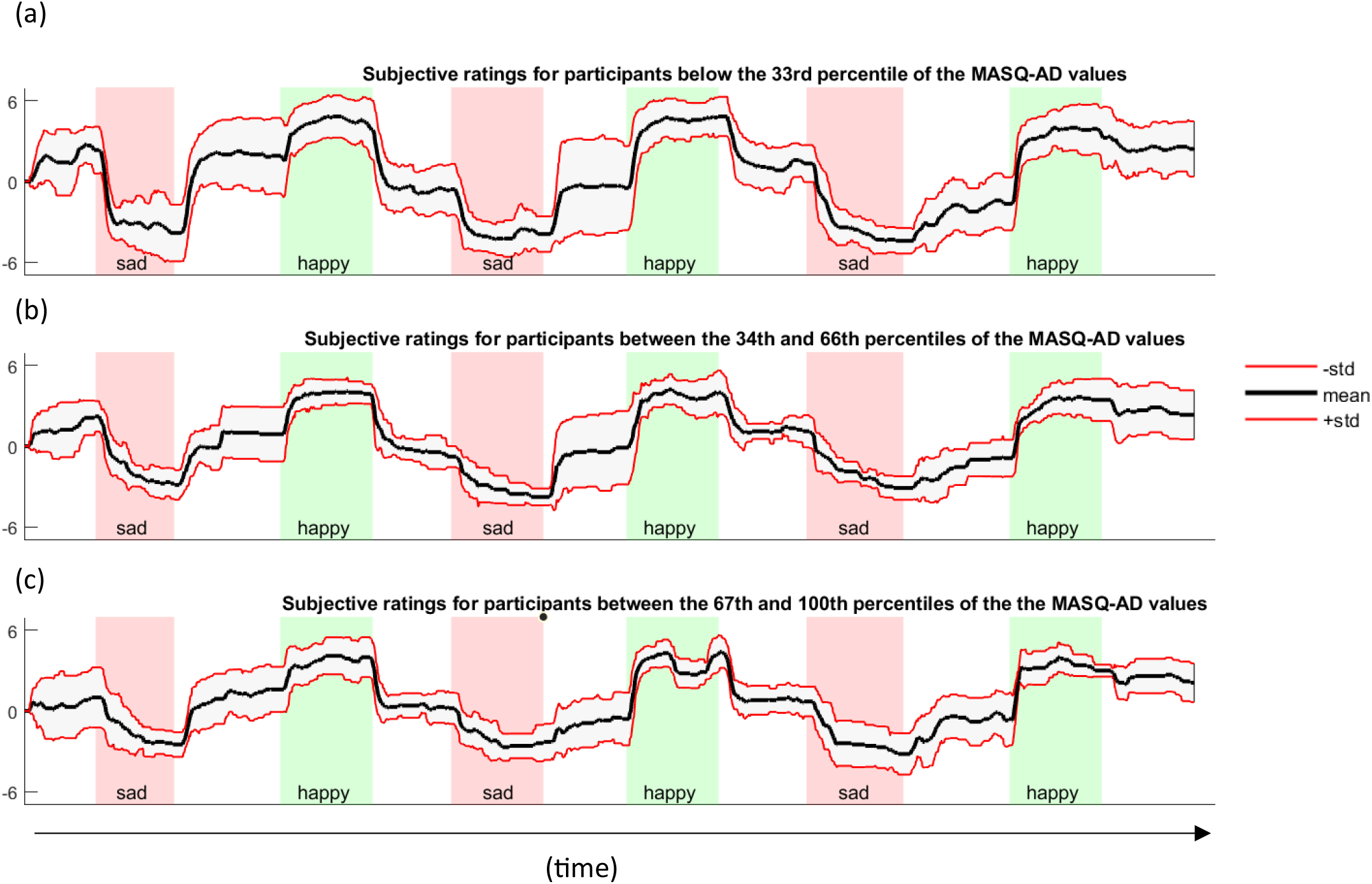
Continuous subjective ratings throughout the entire music task, averaged across the participants with an MASQ-AD score (a) in the bottom 33% of the MASQ-AD values; (b) between the 34^th^ and the 66^th^ percentiles of the MASD-AD values; and (c) in the top 33% of the MASQ-AD values. The bold lines refer to the averaged subjective ratings. The red lines represent the standard deviation associated with the mean subjective ratings.

### LEiDA results

K-means clustering analyses revealed an optimal solution of 9 brain states based on the respective Dunn scores for the range of k-values we used. Figure 4 illustrates the direction and degree to which each of the 105 ROIs’ phases projects onto the leading vector V_1_ of each state (a), and the rendering of each of these states on the cortex (b).

**Figure 4.**
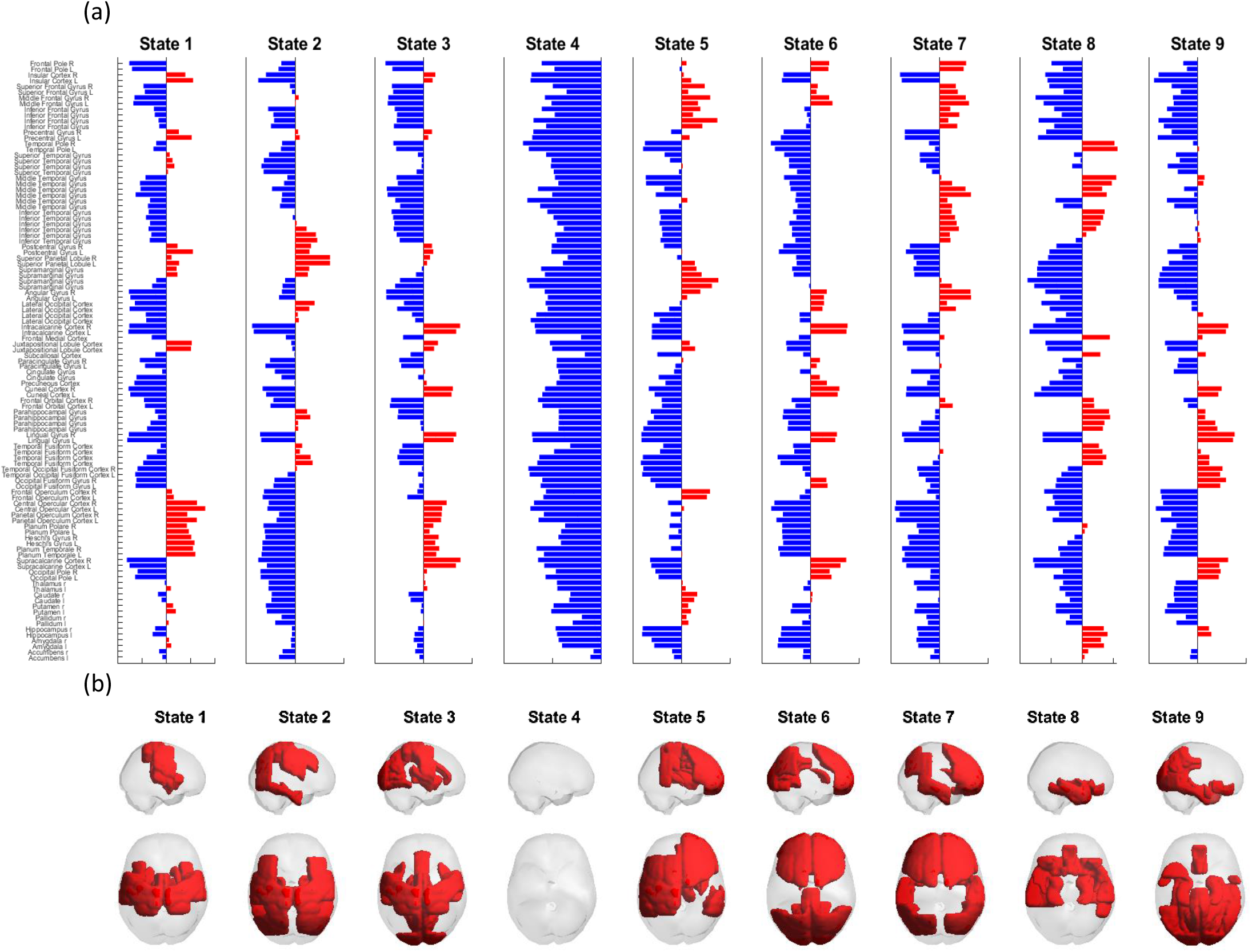
Repertoire of the states deriving from the optimal clustering solution k=9. Top panel (a) refers to the contribution of each brain region to each state. A blue colour represents a negative projection, while a red colour refers to a positive projection. Bottom panel (b) represents the rendering of all brain regions with positive projections onto the leading vector of that state.

### Correlations between each LEiDA state and each Yeo network

State 4, which occurred 14% of the time on average, did not significantly correlate with any Yeo network and can therefore be considered as the global coherence network where all the ROIs’ phases point in the same direction (i.e., negative sign). In contrast, the other eight states are each made up of a specific set of ROIs whose phases have a positive sign (illustrated in red in Figure 4b) and are in synchrony between themselves but out of synchrony with the rest of the ROIs. This limited group of ROIs forms a meaningful network that spatially overlapped with one or more Yeo networks (Lord et al., 2019; Yeo et al., 2011). State 1, which occurred 13% of the time, significantly correlated with the somato-motor and ventral attention networks of Yeo et al. (2011) (r=0.72 and 0.47, respectively); state 2 (9%), with the Dorsal Attention Network (DAN; r=0.58); state 3 (9%), with the visual and somato-motor networks (r=0.43 and 0.42, respectively); state 5 (12%), with the ventral attention and fronto-parietal networks (r=0.47 and 0.53); state 6 (9%), with the visual network (r=0.50); state 7 (11%), with the fronto-parietal and DMN networks (r=0.60 and 0.56, respectively); state 8 (11%), with the limbic network (r=0.60); and state 9 (12%), with the visual network (r=0.73).

### LEiDA metrics across the entire task

#### MASQ-AD and LEiDA metrics

##### Probability of occurrence

A significant positive correlation was identified between the MASQ-AD questionnaire and the probability of occurrence of state 2 (i.e., DAN) (r=0.59, pFDR=0.009; Figure 5). There was no significant correlation between the MASQ-AD and the probability of occurrence of the other states.

**Figure 5.**
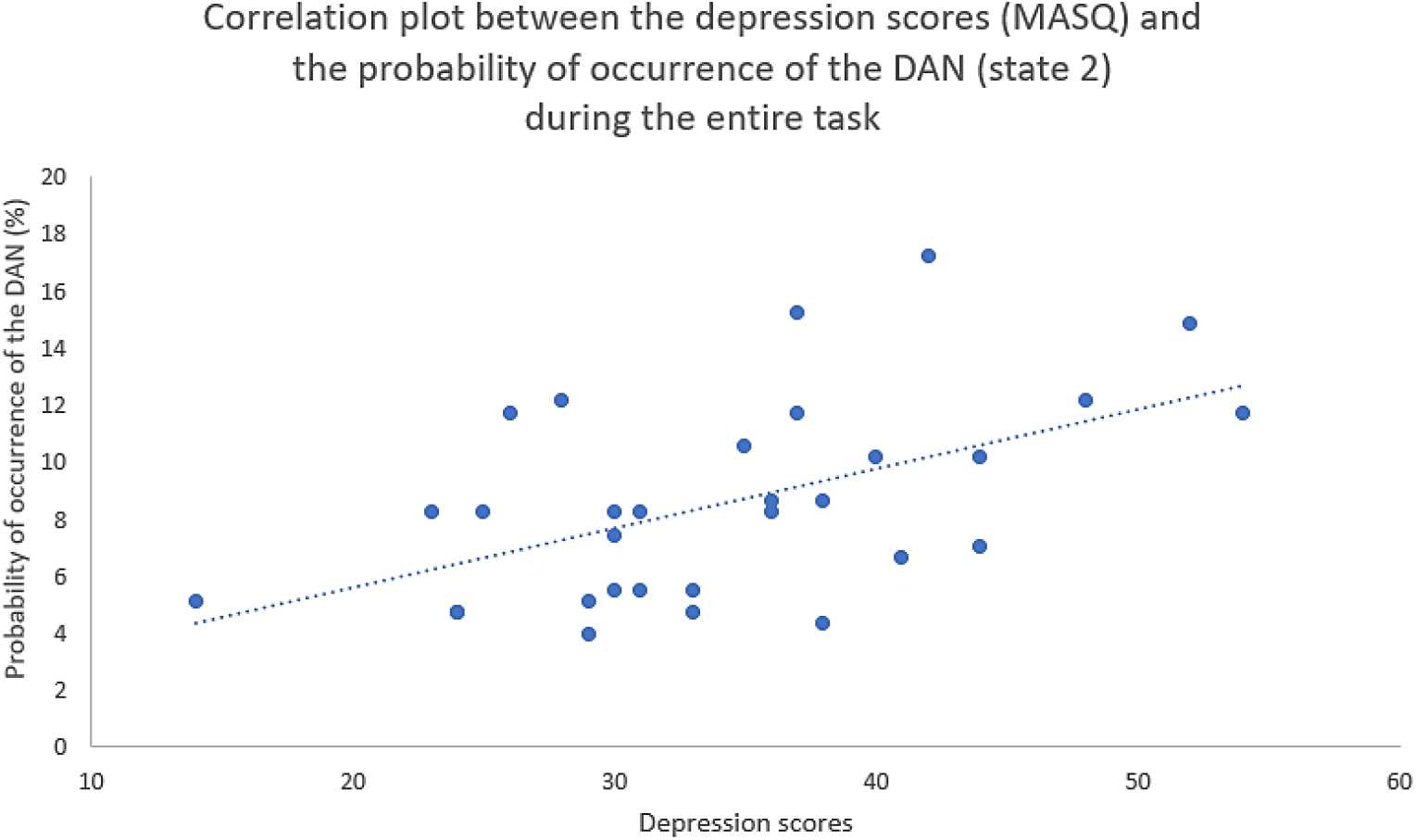
Scatter plot representing the depression scores (MASQ-AD; x axis) and the probability of occurrence of the Dorsal Attention Network (DAN; state 2; y axis), with trendline.

##### Lifetime

There was no significant correlation between the MASQ-AD and the lifetime of any of the states.

##### Switching probability

There was a significant positive correlation between the MASQ-AD and the probability of switching from the global coherence network (state 4) to the DAN (state 2) (r=0.544; p_uncorr_=0.003). We also found a significant positive correlation between the MASQ-AD and the probability of switching from the visual network (state 6) to the ventral attention and fronto-parietal networks (state 5) (r=0.480; p_uncorr_=0.011) and from the fronto-parietal and DMN networks (state 7) to the ventral attention and fronto-parietal networks (state 5) (r=0.384; p_uncorr_=0.048). However, none of these correlations survived FDR correction (pFDR>0.05).

### Emotional blunting/variability and LEiDA metrics

There was a significant negative correlation between the probability of occurrence of the DAN (state 2) and the within-subject STD (i.e., spread in the subjective ratings) (r=-0.393, pFDR=0.043), and a non-significant negative correlation between this same network state (i.e., state 2) and the within-subject RMSSD (r=-0.348, pFDR=0.076) (Figure 6).

**Figure 6.**
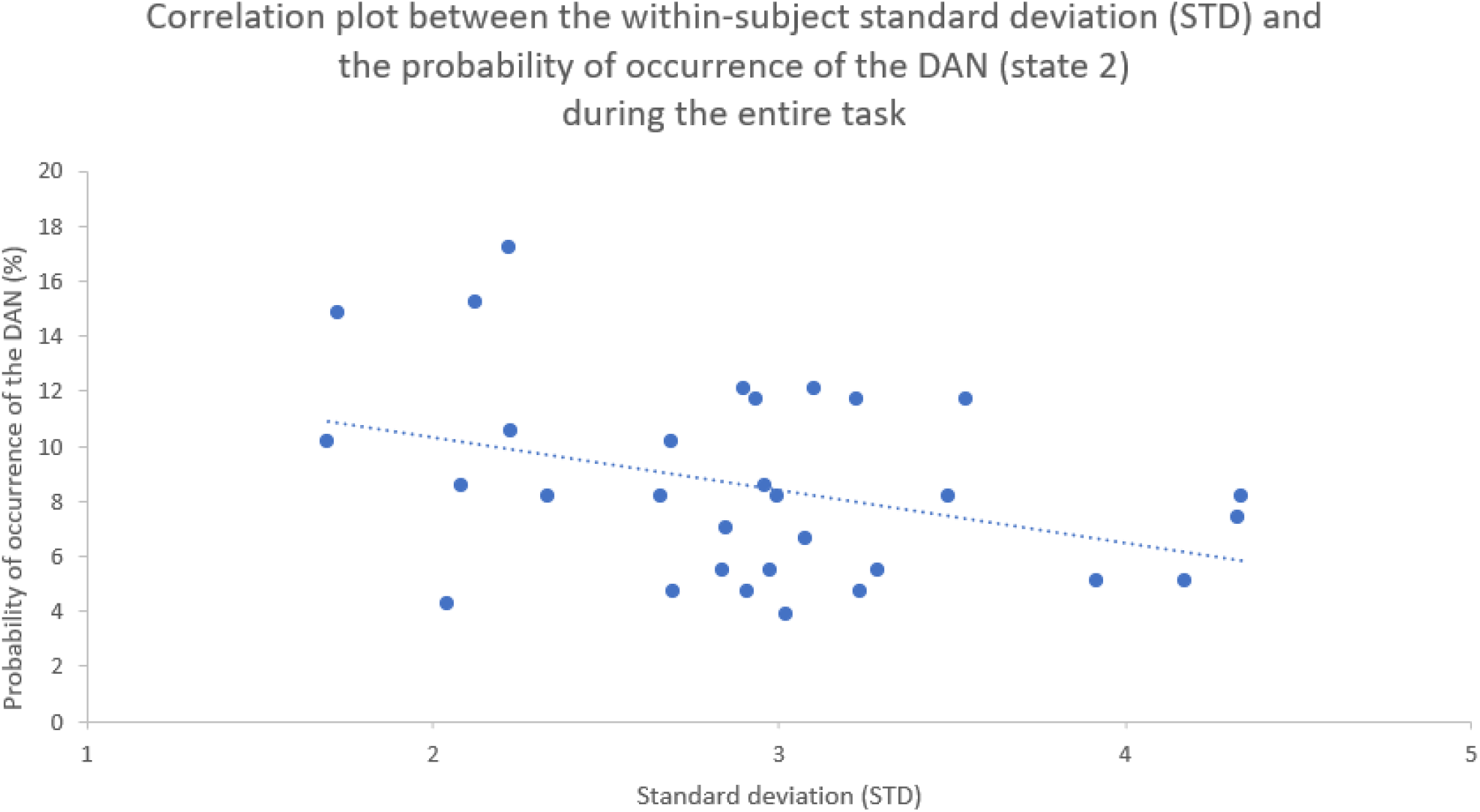
Scatter plot representing the within-subject standard deviation (STD; x axis) and the probability of occurrence of the Dorsal Attention Network (DAN; state 2; y axis), with trendline.

### Mediation

The first step of our mediation analysis showed that the probability of occurrence of the DAN significantly positively predicted anhedonic depressive symptoms (i.e., MASQ-AD; a=0.54, p=0.0012) which, in turn, significantly negatively predicted the within-subject standard deviation of the subjective ratings (i.e., STD; b=-0.62, p=0.0048).

Step 2 revealed a significant indirect effect of the probability of occurrence of the DAN on STD, through MASQ-AD (i.e., ab=-0.33, bootstrapped confidence interval [-0.66 -0.04]). Additionally, the direct effect of the probability of occurrence of the DAN on STD was non-significant (c’=-0.03, p>0.05), while the total effect was significant (c=-0.36, p=0.0425). Taken together, these findings demonstrate that anhedonic depressive symptoms fully mediated the relationship between the probability of occurrence of the DAN and STD (Figure 7).

**Figure 7.**
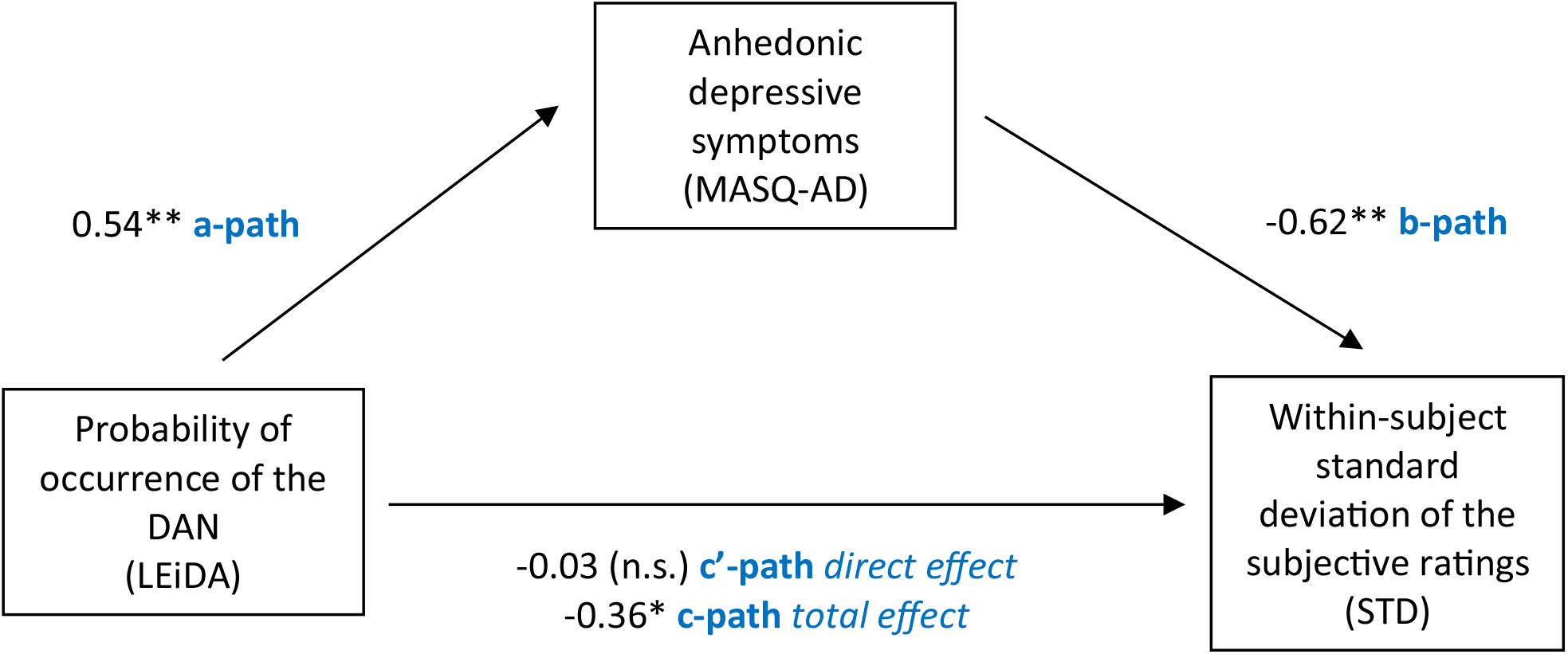
Summary findings of the mediation analysis displaying standardized regression coefficients (betas) and their level of significance for each analysis: *=p<0.05; **=p<0.01; n.s.=non-significant. We found that anhedonic depressive symptoms fully mediated the relationship between the probability of occurrence of the Dorsal Attention Network (i.e., DAN) and the within-subject standard deviation of the subjective ratings (i.e., STD).

### LEiDA metrics with a focus on specific song types

Additional analyses revealed a significant positive correlation between the MASQ-AD and the probability of occurrence of state 2 (i.e., DAN) during neutral songs overall (r=0.606, pFDR<0.001). There was also a significant positive correlation between the MASQ-AD and the probability of occurrence of state 2 (i.e., DAN) during neutral-following-sad songs (r=0.698, pFDR<0.001) and a significant positive correlation between the MASQ-AD and the probability of occurrence of state 2 during neutral-following-happy songs (r=0.459, p_uncorr_=0.016), however the latter did not survive FDR correction.

To illustrate the difference in transition profile from the sad songs to neutral songs in participants with higher levels of depressive symptoms compared to participants with lower levels of depressive symptoms, we split the sample into two groups based on the median of the MASQ-AD scores (34). Figure 8 represents the mean of the probability of occurrence of the sad songs and of the neutral-following-sad songs for the DAN after splitting the participants into two groups.

**Figure 8.**
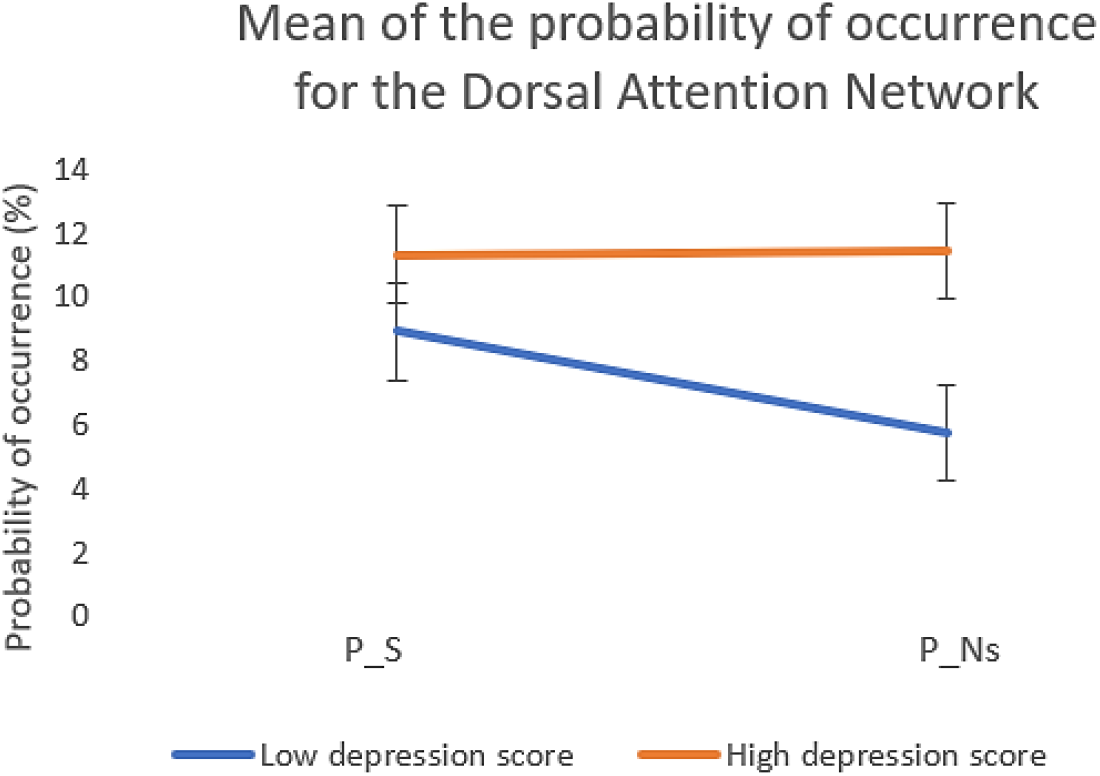
Mean and standard deviation of the probability of occurrence during sad songs (P_S) and during neutral-following-sad songs (P_Ns) for participants low in depression (MASQ-AD score < 34; blue) and high in depression (MASQ-AD score >= 34; orange).

## Discussion

In line with our first hypothesis, we observed an atypical pattern of emotion dynamics (i.e., STD and RMSSD metrics) in participants with higher levels of anhedonic depressive symptoms (i.e., MASQ-AD). In fact, the phenomenon of blunted emotional reactivity previously described by the ECI theory (Rottenberg et al., 2005) aligns with our behavioural findings, showing a negative correlation between MASQ-AD scores and the ‘happy’ beta (i.e., averaged ratings during happy songs), a positive correlation between MASQ-AD scores and the ‘sad’ beta (i.e., averaged ratings during sad songs), and a negative correlation between MASQ-AD scores and both STD and RMSSD scores. However, these findings are inconsistent with the ESM studies reviewed by Houben et al. (2015), where participants were asked to provide information about how they felt every few hours throughout the day, in the absence of any standardized stimulus. Even though the ESM method is ecological, it does not provide information as to whether the emotional responses are adaptive and context-appropriate. Indeed, the higher STD and RMSSD observed in ESM studies in more depressed individuals may be representative of differences in the idiosyncratic environmental contexts that they are typically more exposed to, such as higher levels of bullying and workplace violence and fewer exposure to pleasant events (Eyuboglu et al., 2021; Rottenberg, 2017; Shi et al., 2020).

Partly in line with our second hypothesis, atypical behaviour of the DAN was seen in healthy participants with greater levels of anhedonic depressive symptoms. However, DMN, fronto-parietal and visual network activity patterns were unrelated to anhedonia. For the DAN, those reporting greater levels of anhedonia exhibited a higher probability of the network’s occurrence throughout the task. In fact, attentional difficulties have previously been observed in clinical depression (see Rock et al., 2014 for a meta-analysis). The DAN is involved in externally oriented attention, and atypical behaviours of this network have been observed both at rest (see Kaiser et al., 2015 for a meta-analysis) and during task performance (Sambataro et al., 2017) in MDD. In the context of LEiDA, previous studies found a lower engagement of the DAN in subclinical depression (Alonso-Martínez et al., 2020), which contrasts with our findings. However, our study required the participants to actively engage in a task by providing continuous subjective ratings, whereas Alonso-Martínez et al. (2020) measured the behaviour of this network at rest, which could explain the discrepancy between findings. In fact, in resting-state studies, the absence of any manipulation of attention or emotion hinders the drawing of conclusions about the relationship between atypical network recruitment and self-reported emotional difficulties (Poldrack, 2006). Indeed, it is difficult to infer specific patterns of affective processes from unconstrained brain activity (Finn, 2021).

We also demonstrated, for the first time to our knowledge, that blunted emotional responses (i.e., lower within-subject STD of the subjective ratings) were associated with an increased probability of occurrence of the DAN throughout the task. This is at least partly consistent with our third hypothesis. In fact, further analyses revealed that MASQ-AD fully mediated the relationship between DAN recruitment and emotional blunting. More specifically, DAN recruitment positively predicted anhedonic depressive symptoms which, in turn, negatively predicted the magnitude of the subjective ratings (i.e., within-subject standard deviation). These findings are consistent with the notions of numbing out and atypical engagement with the external environment previously described by Rottenberg (2017) and the Constructionist theory (Barrett, 2017). Indeed, according to Constructionism, the depressed brain keeps making incorrect predictions about the body’s energy needs as if it was chronically predicting and reliving painful events from the past when the metabolic needs were high and costly (Clark-Polner, Wager, Satpute & Barrett, 2016). As such, the depressed brain pays more attention to its surroundings, ready to fight off and meet energy needs that are non-existent in the present. This leads to the entire system eventually shutting down in an attempt to cut “spending”, which translates into emotional blunting and disengagement (Rottenberg, 2017).

In further exploratory analyses, we also found that the relationship between the MASQ-AD scores and DAN activity was even more evident while participants listened to neutral songs, particularly neutral music that followed sad music. The recruitment of the DAN remained elevated in the transition from emotional to neutral pieces of music in more depressed individuals, while it decreased for less depressed participants. Together with the lower STD scores, these findings speak in favour of the ECI theory (Rottenberg et al., 2005) which describes a lack of sensitivity in the transition from one type of emotional event to the next. This could reflect a carry-over effect as observed in previous studies where more depressed individuals struggle to recover from negative emotional states (Ebrahimi et al., 2021; Minaeva et al., 2021), ). In fact, larger within-subject levels of lethargy that carried over across days have previously been found to temporally predict increases in within-subject levels of anhedonia on the consecutive day (Ebrahimi et al., 2021). This difficulty to recover from a negative emotional state has also been described in the context of shifting impairments in the transition from negative to neutral information in dysphoric adolescents (Wante et al., 2017).

### Limitations

#### This study has several limitations

Firstly, “neutral” songs may have affective connotations. Indeed, even though in our study averaged ratings were significantly different between happy and neutral songs, and between sad and neutral songs, previous studies have shown that neutral stimuli may sometimes be interpreted as slightly negative (Lee et al., 2008), and this might have affected our findings.

Second, this study did not control for the perceptual properties of each song. Indeed, it is well-known that emotional responses to music may vary with features such as loudness, pitch level or sharpness (Coutinho & Cangelosi, 2011). Future studies may investigate whether the relationship between the severity of anhedonic depressive symptoms, the subjective ratings and attentional networks recruitment was somehow related to some of the song features. Indeed, this may answer questions about the interactions between attention, perception and emotions.

Thirdly, overall, the happy songs were rated as more familiar by the participants than the sad songs. Some happy songs may have been associated with specific memories in some participants, which might have been reflected in their ratings. However, we did control for familiarity ratings in all analyses. Future studies should consider a post-scan debriefing session with the participants to further investigate potential autobiographical memories at play during the experiment. Additionally, we asked the participants to listen to the pieces of music outside of the scanner once before fMRI data acquisition. Even though this allowed us to successfully collect information about the participants’ familiarity with the different pieces of music, it might also have created a repetition effect, potentially impacting the emotional responses to the music whilst inside the scanner (Margulis, 2013).

In addition, this study was carried out in a healthy sample, which means that, at this stage, we cannot generalise these findings to clinical populations, but this encourages future research to explore dynamic responses to naturalistic paradigms in clinical samples, such as MDD.

Finally, it is worth noting that the background noise generated by the scanner may have impacted on the participants’ emotional responses to the different pieces of music. Indeed, previous research has shown that scanner noise may distort affective brain processes (Skouras et al., 2013). Future studies would benefit from exploring affective processing in the context of quieter MRI sequences such as Looping Star (Wiesinger, Menini & Solana, 2019).

## Conclusion

In this study, we explored behavioural and brain network dynamics as a function of anhedonic depressive symptoms severity in healthy adults during an emotionally provocative music listening task. Our findings highlight an atypical behaviour of the Dorsal Attention Network in healthy participants with higher levels of anhedonic depressive symptoms, which was associated with a sustained blunted emotional response to both happy and sad songs. The elevated recruitment of the Dorsal Attention Network during emotional pieces of music carried over into subsequent affectively neutral music. Future research should explore whether these findings could be generalised to a clinical population (i.e., Major Depressive Disorder).

