## Supplementary material for "Anhedonia severity mediates the relationship between attentional networks recruitment and emotional blunting during music listening"

### Supplemental Material

#### Supplementary Material 1a

*Personality and mental health questionnaires: materials and methods*

*Big Five Inventory (BFI) – neuroticism subscale*

We used the 8-item neuroticism subscale of the 44-item BFI (John & Srivastava, 1999) to measure neuroticism levels. The subscale is composed of items that query to what extent participants typically agree with statements such as "I see myself as someone who is depressed, blue". Agreement with items was rated on a 7-point scale ranging between 1 (e.g., strongly disagree) and 7 (e.g., strongly agree). The neuroticism subscale has previously shown good internal consistency ($\alpha$=.82) and excellent test-retest reliability ($\alpha$=.93) in a university sample (Arterberry et al., 2014). Overall scores range between 8 and 56, with higher scores indicating higher levels of neuroticism.

*Rumination-reflection questionnaire (RRQ) – rumination subscale*

The rumination subscale of the RRQ (Trapnell & Campbell, 1999) is made up of 8 items such as "Long after an argument or disagreement is over with, my thoughts keep going back to what happened". Participants rated each statement on a five-point scale ranging between 1 (e.g., strongly disagree) and 5 (e.g., strongly agree) in terms of how they typically applied to them. Overall scores range between 5 and 40, and higher scores reflect higher rumination levels. The RRQ has demonstrated excellent internal consistency alpha in university samples ($\alpha$=.91; Garrido, 2018).

*Perth Emotional Reactivity Scale – short version (PERS-short)*

The short version of the PERS (Preece, Becerra & Campitelli, 2019) is composed of 18 items that inquire about how participants typically respond to emotional events. For each item, participants are required to indicate how closely the statement typically applies to them on a five-point scale ranging between 1 (very unlike me) and 5 (very like me). The PERS-short assesses emotional reactivity for positive and negative emotions separately across three dimensions: activation (e.g., "I tend to get happy very easily" or "I tend to get pessimistic about negative things very quickly"), intensity (e.g., "I experience positive mood very strongly" or "My negative feelings feel very intense") and duration (e.g., "When I'm happy, the feeling stays with me for quite a while" or "When I'm upset, it takes me quite a while to snap out of it"). The scale allows for the calculation of a general positive reactivity (GPR) score obtained by summing up all the items in the positive activation, intensity and duration subscales; and a general negative reactivity (GNR) score by summing up items from the negative activation, intensity and duration subscales. The GPR and the GNR scales have been shown to have excellent internal consistency ($\alpha$=.92 and $\alpha$=.91, respectively) in a large community sample (Preece, Becerra & Campitelli, 2019).

#### Supplementary Material 1b

*Personality and mental health questionnaires: behavioural results*

The mean and between-subject standard deviation for all personality and mental health questionnaires can be found in Supplementary Material Table 1. Correlations between each pair of questionnaires, and associated p-values, are presented in Supplementary Material Table 2.
Because the focus of this study was on depressive symptoms, and because the MASQ-AD was highly correlated with the other 4 questionnaires, we chose to focus on the MASQ-AD questionnaire for the rest of the analyses.

| Questionnaires (scales) | Mean | SD |
|---|---|---|
| MASQ-AD (anhedonic depression) | 34.26 | 8.843 |
| BFI (neuroticism) | 27.48 | 8.640 |
| RRQ (rumination) | 25.74 | 5.633 |
| PERS (GPR) | 33.52 | 5.189 |
| PERS (GNR) | 26.42 | 7.915 |

**Supplementary Material Table 1. Descriptive statistics providing the mean and standard deviation for each questionnaire. SD=standard deviation.**

| | MASQ-AD | Neuroticism | Rumination | GPR | GNR |
|---|---|---|---|---|---|
| **MASQ-AD** | | 0.527 (0.002) | 0.377 (0.037) | -0.628 (<0.001) | 0.358 (0.048) |
| **Neuroticism** | | | 0.657 (<0.001) | -0.383 (0.034) | 0.829 (<0.001) |
| **Rumination** | | | | -0.272 (0.138) | 0.598 (<0.001) |
| **GPR** | | | | | -0.114 (0.541) |
| **GNR** | | | | | |

**Supplementary Material Table 2. Correlations and p-values (in parentheses) between pairs of questionnaires. MASQ-AD= Mood and Anxiety Symptoms Questionnaire; GNR=General Negative Reactivity subscale of the Perth Emotional Reactivity Scale; GPR=General Positive Reactivity subscale of the Perth Emotional Reactivity Scale.**

**Supplementary Material 2a**

*Familiarity ratings and music background: behavioural analysis*

Because controlling for the familiarity ratings of each song would have added 13 covariates, thus significantly lowering the number of degrees of freedom in our analyses, we first calculated the mean of the familiarity ratings for each song type by averaging across the familiarity ratings of all songs of the same type. This led to three summary measures: Familiarity_H (i.e., familiarity with the happy songs), Familiary_S (i.e., familiarity with the sad songs), Familiarity_N (i.e., familiarity with the neutral songs).
The second step consisted of identifying the music background question, of the four answered by participants, that best explained the variance in the familiarity ratings across all song types, and which we would then use as a nuisance covariate for subsequent analyses. We computed the Pearson's correlations between each familiarity summary measure and each of the four music background questions, as well as the associated coefficients of determination representing the percentage of variance in the familiarity ratings explained by each music background question. As question 3 (i.e., I

have a broad knowledge of classical music) consistently exhibited the highest coefficients of determination and correlation coefficients for the four familiarity summary measures, the question 3 scores were added to gender and age as a nuisance covariate for the subsequent analyses.

**Supplementary Material 2b**

*Familiarity ratings and music background: behavioural results*

The mean and standard deviation for each music background question and for familiarity ratings for each song type can be found in Supplementary Material Table 3 and Supplementary Material Table 4.

| Familiarity ratings per song type | Mean | SD |
|---|---|---|
| Familiarity_H | 2.290 | 1.006 |
| Familiarity_S | 1.387 | 0.615 |
| Familiarity_N | 1.823 | 0.781 |

**Supplementary Material Table 3. Mean and between-subject standard deviation of the familiarity for each song type. The mean of the familiarity ratings for each song type was calculated by averaging across the familiarity ratings of all three songs of the same type. This led to three summary measures: Familiarity_H (i.e., familiarity with the happy songs), Familiarity_S (i.e., familiarity with the sad songs) and Familiarity_N (i.e., familiarity with the neutral songs). SD= standard deviation of the scores across participants for each familiarity type.**

| Music background | Mean | SD |
|---|---|---|
| Question 1: “I enjoy listening to music” | 4.585 | 0.916 |
| Question 2: “I enjoy listening to classical music” | 2.945 | 1.188 |
| Question 3: “I have a broad knowledge of classical music” | 1.970 | 0.984 |
| Question 4: “I can play a musical instrument” | 2.704 | 1.452 |

**Supplementary Material Table 4. Descriptive statistics representing the mean and standard deviation for each music background question. Participants were required to rate each question in terms of how much it applied to them. For each question, the scores ranged between 1 and 5. A score of 1 meant that the statement did not apply to them at all, while a score of 5 meant that it extremely applied to them. SD = standard deviation of the scores across participants for each question.**

The coefficients of determination and correlation coefficients between the music background questions and the familiarity scores can be found in Supplementary Material Table 5 and Supplementary Material Figure 1.

| | Familiarity_happy | | Familiarity_sad | | Familiarity_neutral | |
|---|---|---|---|---|---|---|
| | CC | CD | CC | CD | CC | CD |
| Question 1 | -0.022 (0.907) | 0 | -0.048 (0.800) | 0.002 | 0.138 (0.460) | 0.019 |
| Question 2 | 0.224 (0.226) | 0.05 | 0.183 (0.324) | 0.033 | 0.458 (0.010) | 0.21 |
| Question 3 | 0.431 (0.016) | 0.186 | 0.379 (0.035) | 0.143 | 0.542 (0.002) | 0.294 |

| | | | | | | |
|---|---|---|---|---|---|---|
| Question 4 | 0.320 (0.079) | 0.102 | 0.186 (0.316) | 0.0346 | 0.270 (0.142) | 0.0729 |

**Supplementary Material Table 5. Correlation coefficients (and p-values) and associated coefficients of determination between each item of the music background question and each of the family ratings per song type. The higher the CC and the CD, the greater the relationship between the question and the familiarity for the particular song type.**

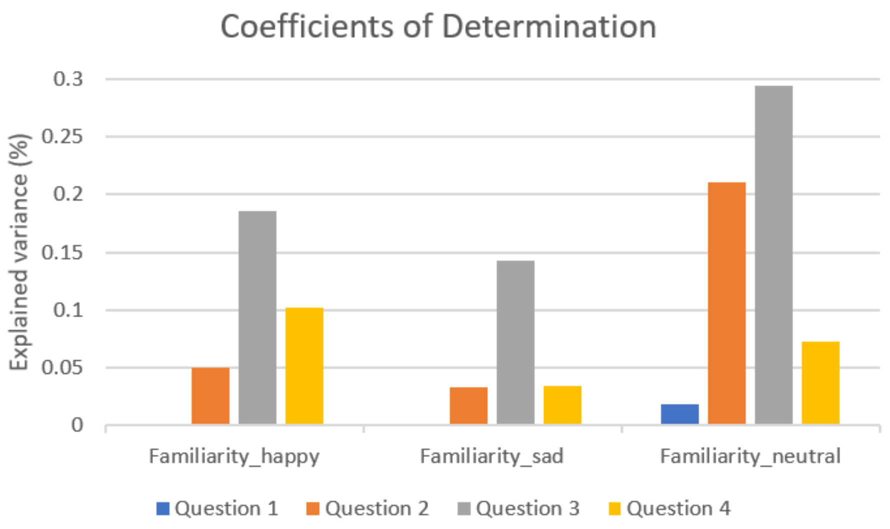


**Supplementary Material Figure 1. Coefficients of Determination representing the percentage of variance in familiarity ratings explained by each music background question. Question 1 = "I enjoy listening to music"; question 2 = "I enjoy listening to classical music"; question 3 = "I have a broad knowledge of classical music"; question 4 = "I can play a musical instrument".**

**Supplementary Material 3a**

*MASQ-AD and mean subjective ratings: behavioural analysis*

For each participant, the mean subjective ratings for a given song type were estimated by first building a Generalized Linear Model (GLM) with a constant/intercept and regressors for each song type. Each regressor encoded the time (i.e., onset and duration) that each song of that song type was playing. For each participant, we thus obtained three beta values (i.e., 'happy', 'sad' and 'neutral').

To evaluate whether the mean subjective ratings differed significantly across all three song types, we first carried out a repeated-measures ANOVA, followed by two post-hoc paired t-tests to explore further whether there was a significant difference between 'happy' and 'neutral' betas, and between 'sad' and 'neutral' betas.

To identify whether there was an association between the anhedonic depression symptoms and the average subjective ratings attributed to each song type, we then correlated the 'happy', 'sad' and 'neutral' betas with the MASQ-AD scores, controlling for age, gender and music familiarity ratings corresponding to the relevant beta valence.

**Supplementary Material 3b**

*MASQ-AD and mean subjective ratings: behavioural results*

A repeated-measures ANOVA revealed that the mean subjective ratings differed significantly across happy, sad and neutral songs ($F(2,52)=6.401$, $p=0.003$). Post-hoc paired t-tests showed that the mean subjective ratings were significantly higher for happy compared to neutral songs ($t(30)=14.536$ ($p<0.001$) and significantly lower for sad compared to neutral songs ($t(30)=-11.778$, $p<0.001$).

There was a significant negative correlation between MASQ-AD and the 'happy' beta ($r=-0.41$, $p_{FDR}=0.03$), and a significant positive correlation between MASQ-AD and the 'sad' beta ($r=0.61$, $p_{FDR}=0.0014$). There was no significant correlation between MASQ-AD and the 'neutral' beta ($p>0.05$).
